# Matrix Stiffness Regulates Liver Sinusoidal Endothelial Cell Function Mimicking Responses in Fatty Liver Disease

**DOI:** 10.1101/2020.01.27.921353

**Authors:** Vaishaali Natarajan, Michael Moeller, Carol A. Casey, Edward N. Harris, Srivatsan Kidambi

## Abstract

Liver sinusoidal endothelial cells (LSECs) are a highly specialized endothelial cell that participates in numerous liver metabolic activities and collectively function as a scavenger system in the liver by removing waste macromolecules playing a vital role in the balance of lipids, cholesterol, and vitamins. Prior to hepatic fibrosis, independent of their etiology, LSECs become highly pro-inflammatory, capillarized (loss in fenestrations), and loss in specialized receptors (Stabilin-1, Stabilin-2, CD31 and SE-1). Liver fibrosis also leads to significant loss in the endocytosis function of LSECs. Thus understanding regulation of LSEC phenotype may be critical to understanding fibrosis. Extensive remodeling of the extracellular matrix occurs during fibrosis that leads to liver stiffening. The role of matrix stiffness as related to subtle but pivotal changes in LSECs physiology is under explored. The **overall goal** of our study is the development and implementation of a platform that enables the convergence of engineered cell microenvironments with the phenotypic and functional analysis of LSECs. Using our innovative biomimetic liver fibrosis model that allows modulation of substrate stiffness, we investigated the role of liver matrix stiffness in modulating LSECs function in fibrotic-like microenvironment. Primary LSECs were cultured on our novel polymer film coated polydimethylsiloxane (PDMS) gels with 2 kPa, 9 kPa 25 kPa and 55 kPa elastic modulus mimicking healthy, early fibrotic, fibrotic and extremely fibrotic substrates. SEM was used to image to fenestrations of LSECs and HA endocytosis assay was performed to measure the LSECs function. LSECs cultured on stiffer environment had significant remodeling of cytoskeletal proteins and morphology indicated of stress fibers. Also we observed that LSECs on fibrotic substrates resulted in loss of fenestrations (capillarization). This is critical as capillarization has been show to precede hepatic fibrosis and “capillarized” LSECs lose the ability to promote hepatic stellate cell (HSC) quiescence. LSECs on stiffer environment also had higher expression of cell adhesion molecules, VCAM-1 and ICAM-1 indicating the loss of phenotype of the cells. Fibrotic stiffness also impeded the HA endocytosis in LSECs, one of the main functions of the cells. These data suggest a plausible mechanism that increased stiffness modulates hepatocyte and LSEC function causing liver functional failure. Similar effect was observed in LSECs isolated from Non-Alcoholic Fatty Liver Disease (NAFLD) rat models indicating correlation to physiological conditions. Together, all these data demonstrates the plausible role of stiffness in regulating LSECs function and contribute to HSC activation and progression of liver fibrosis.

## 1. Introduction

Liver sinusoidal endothelial cells (LSECs) are the highly specialized cell type lining the rich network of capillaries/sinusoid that regulate the blood flow to hepatic cords (1). Unlike vascular endothelial cells, LSECs possess unique morphological features such as lack of basement membrane and transcellular fenestrations that enable the cells to act as a dynamic filter by regulating the transport of solutes and macromolecules between the lumen of the liver and the parenchyma and, excluding the transport of larger particles such as cells and chylomicrons (2). Additionally, the presence of a high density of endocytic vesicles in these cells allows for scavenging action and clearance of unwanted molecules from the blood (3). Maintenance of the morphological integrity is key in the maintenance of the complex repertoire of LSEC function and consequentially, capillarization/dedifferentiation of LSECs is evaluated by the loss in fenestrations, decreased endocytic capacity and appearance of basement membrane (4). Studies report that capillarization event of the highly specialized LSEC phenotype is a vital gatekeeper event in the progression of metabolic liver diseases such as hepatitis (5) nonalcoholic fatty liver disease (6), alcoholic liver disease (7) (8) and liver fibrosis (9) (10). Despite extensive research supporting the importance of LSECs in liver pathophysiology, the mechanistic aspects of LSEC capillarization are poorly established.

Chronic stress to the liver due to etiologies such as viral infections, alcohol abuse, and non-alcoholic steatosis can result in liver fibrosis which is accompanied by uncontrolled deposition, remodeling, and abnormal turnover of extracellular matrix (ECM) proteins (11). Progression of fibrosis relies on the interplay of the liver microenvironmental (LME) elements such as heterotypic cell-cell interactions, cell-ECM interactions, chemical signals and mechanical factors (12). The hallmark of liver fibrosis is the increase in mechanical stiffness of the tissue due to the disrupted balance between ECM production and renewal (11). Increased stiffness of the liver is the most universally recognized clinical diagnostic marker for establishing various stages of liver fibrosis (13, 14). The unchecked increase in the LME stiffness due to excessive accumulation of ECM proteins results in a further interruption in cell-cell communication and ultimately paves the way for scar tissue taking over the liver resulting in irreversible cirrhosis. Cirrhosis is a leading cause of death since cirrhotic liver subsequently causes fatal hepatic conditions such as hepatocellular carcinoma, hepatic encephalopathy, renal failure and systemic toxicity (11, 15).

Due to their viscoelastic nature, adherent cells of several origins demonstrate a mechanosensitive behavior when subjected to changes in the mechanical microenvironment of the surrounding (16). In the liver, previous studies have established this mechanosensitivity in hepatocytes (17) (18) (19) and hepatic stellate cells (20) but only two such studies have been conducted on LSECs (21, 22). Research suggests that capillarization of LSECs is a permissive event for liver fibrosis and, triggers a series of pro-fibrotic responses in the liver that is mediated through activation of hepatic stellate cells. Therefore, identifying the various LME triggers that promote LSEC capillarization is a vital step towards targeting LSECs in anti-fibrosis interventions. We hypothesize that varying the stiffness of the organ plays a pivotal role in the progression of liver fibrosis and considering the important role of LSECs in this liver disease; there is a critical need to establish the correlation between the mechanical stiffness in the liver and the LSEC phenotype.

A vast portion of LSEC research relies on animal models due to the lack of robust *in vitro* culture systems for these cell types. Despite their physiological relevance, animal models form an inherently complex tool for correlating phenotypic changes of a particular cell type with the LME elements, without considering exogenous factors. Hence, designing *in vitro* models to capture and elucidate the changes in the LSEC phenotype, corresponding to the healthy and fibrotic milieu, will prove to be an invaluable tool towards the study of liver fibrosis.

Here we report a novel *in vitro* model to investigate the changes in LSEC phenotype that are driven solely by the drastic change in the mechanical environment of the liver at healthy and advanced fibrosis stages. We created the *in vitro* platform by employing two different precursors of polydimethylsiloxane (PDMS), namely Sylgard 527 and Sylgard 184 to create substrates with tunable stiffness (23). The different weight-based blending of the two PDMS precursors was used to create culture platforms mimicking “healthy” (soft substrate) and “fibrotic” (stiff substrate) liver. Our platform enables investigation of the isolated effect of mechanical changes in liver microenvironment on LSECs, without varying the surface ligand density or topographical features. Our results suggest that LSECs cultured on the “stiff” substrate demonstrated rapid capillarization, as witnessed by visualization of fenestrations, cell adhesion marker expression analysis, and cytoskeletal investigation. We also observed that LSECs undergo a loss in HA endocytosis function in a fibrosis-like culture environment. On the other hand, the healthy liver mimic sustained the normal LSEC phenotype for about 48 hours, *in vitro*. Our *in vitro* platforms can serve to recreate LSEC physiology of the healthy and the fibrotic liver and hence, can potentially be used as a tool to understand the complex mechanistic aspects regulating these cells and their role in disease progression.

## 2. Materials and Methods

### 2.1 PDMS Substrate Preparation

*In vitro* healthy and fibrotic liver mimics were created using PDMS based substrates (19). In a technique adapted from Palchesko and coworkers, Sylgard 527 and Sylgard 184 were taken in a specific weight ratio as mentioned in Table. 1. The crosslinking process was carried out using manufacturer’s guidelines. Briefly, for Sylgard 527, equal parts of component A and B were mixed well. Sylgard 184 was mixed in a ratio of 10:1 of elastomer to cross-linking agent. The two precursors were then blended in the desired weight ratio (empirically determined) and poured into 12 well tissue culture plates and cross-linking was carried out at 65 °C.

**Table 1:** The mass percentage of Sylgard 184 and the corresponding stiffness (Young’s modulus) of the substrates

Following overnight cross-linking, the 12 well plates were subjected to oxygen plasma treatment for a 7-minute duration to render the PDMS surfaces hydrophilic (Plasma Cleaner, PDC001, Harrick Plasma, Ithaca NY). Plates were then coated with 0.1 mg/ml rat tail type I collagen solution (in 0.01N acetic acid) overnight. Plates were washed with phosphate buffer saline (PBS) three times and sterilized under UV and used immediately for the culture of LSECs.

### 2.2 Elastic Modulus Determination for the PDMS Substrates

Young’s moduli for soft and stiff PDMS substrates were determined using the TMS-Pro texture analyzer instrument (Food Technology Corporation, Sterling, VA). The height and the diameter of the PDMS samples prepared were determined using Vernier calipers. Following this, the PDMS samples were subjected to compression during which, the force applied and the corresponding displacement achieved, were recorded. The range of force and displacement values were utilized to construct stress-strain curves and the Young’s Modulus for each sample was determined from the linear region of the stress-strain curve.

### 2.3 Collagen Quantification on PDMS Substrate Surface

*Fluorescent Imaging*: 12 well plates containing the PDMS substrates were subjected to plasma treatment for 7 minutes followed by incubation with collagen-FITC for 1 hour at 37 °C. The plates were washed extensively after incubation to get rid of loosely bound collagen-FITC and were viewed/imaged using a fluorescence microscope (Axiovert 40 CFL, Zeiss, Germany).

*BCA quantification:* To quantify the amount of collagen bound per unit surface area on each substrate, collagen deposition was carried out as described in section 2.1 and then RIPA buffer was used to scrape it off the substrate and collected in microfuge tubes. The amount of collagen was determined using Pierce™ BCA Protein Assay Kit (Fisher Scientific, PA). Standard curve was constructed using known concentrations of collagen type-1.

### 2.3 Primary Liver Sinusoidal Endothelial Cell Isolation and Culture

Primary LSECs were isolated from male Sprague-Dawley rats weighing 150-200 grams using a protocol adapted from P.O Seglen (24). All animal procedures were performed in accordance with the IACUC guidelines of the University of Nebraska-Lincoln. All the hepatic cells were isolated from the rat liver using a two-step collagenase digestion procedure. The cell suspension was spun at 150 *g* for 3 minutes to pellet the hepatocytes. Supernatant was spun at 200 g for 10 minutes to pellet the viable non-parenchymal cells. The non-parenchymal cells were suspended in RPMI-BSA and loaded onto a 25%/50% Percoll gradient and spun at 900 g for 20 minutes. The interface at the 25% and 50% gradient was separated and spun again at 200 g for 10 minutes. Following this, the cells obtained from the pellet were suspended in pre-warmed RPMI containing 100 U/ml Penicillin and Streptomycin and placed in an acid washed crystallizing dish in the tissue culture incubator for 15 minutes. The supernatant was collected and spun down at 200 g to obtain the LSECs rich pellet. Cell viability at the end of the process was determined using Trypan blue exclusion test.

The cells were seeded onto the collagen coated PDMS substrates at a seeding density of 400,000/ sq. cm. LSECs culture media was prepared with high glucose DMEM supplemented with 10% FBS, 1 µg/ml VEGF, 0.5 U/ml insulin, 20 ng/ml epidermal growth factor (EGF), 7 ng/ml glucagon, 7.5 mg/ml hydrocortisone and 1% penicillin/streptomycin. Cells were maintained in a humidified 5% CO_2_ incubator at 37 °C.

### 2.4 LSEC Morphology Analysis

Phase contrast images of LSECs cultured on soft and stiff substrates were taken at different time points using an inverted microscope (Axiovert 40 CFL, Zeiss, Germany). In order to visualize the cell attachment, cells were fixed after 6 hours in culture using 4% paraformaldehyde prepared in sterile PBS, permeabilized using 0.1% Triton X 100 and treated with DAPI stain at a concentration of 10 µM. Substrates were imaged using a fluorescent microscope (Axiovert 40 CFL, Zeiss, Germany) under a UV filter.

*Actin Staining:* LSECs were fixed with 4% paraformaldehyde for 15 minutes, followed by permeabilization for 5 minutes using 0.2% Triton X 100 solution prepared in PBS. The cells were rinsed and treated with Alexa Fluor 488 Phalloidin for 30 minutes (Molecular Probes, US). Actin filaments in the cells were imaged using a fluorescent microscope.

### 2.5 Protein Expression

Cells were washed with PBS and lysed in 12 well plates containing the PDMS substrates using 75 µl RIPA buffer (100mM Tris, 5mM EDTA, 5% NP40) supplemented with 1X protease inhibitor cocktail and phenylmethylsulfonyl fluoride (PMSF) by incubating on ice for ten minutes, followed by collection of cell lysates in microfuge tubes. Cell debris was pelleted out and supernatants with proteins were stored away at −80 °C until use. Protein concentration was determined colorimetrically using Pierce™ BCA Protein Assay Kit (Fisher Scientific, PA). Protein was loaded onto SDS-containing polyacrylamide gels (10% acrylamide for VCAM-1/ICAM-1 blots and 5% acrylamide for Stabilin-2 blots) and after PAGE, were transferred onto Immobilon CL membrane (Millipore, MA). Membranes were blocked using 5% skimmed milk for 2 hours at room temperature (RT) following which the blots were incubated overnight at 4 °C in anti-VCAM-1 (Santa Cruz, CA), anti-ICAM-1 (Santa Cruz, CA), anti-stabilin-2 (Santa Cruz, CA) or anti-GAPDH (Millipore, MA) antibodies. Following the primary antibody incubation, the blots were incubated for one hour at RT in near infrared 680nm and 800nm secondary antibody (Fisher Scientific, PA) and signal for protein expression was detected using Odyssey Infrared Imaging System (Li-COR Biosciences, USA). Densitometric analysis of the blots was performed using the Image Studio software associated with Odyssey Imaging System.

### 2.6 Real Time Gene Expression

Total RNA from the cell samples was isolated using Trizol reagent (Life Technologies, USA) following the manufacturer’s protocol. Briefly, the cells were lysed with Trizol for 15 minutes, followed by spinning down at 12,000 g to remove cell debris. The Trizol cell lysate was mixed with one fifth volume choloform and the samples were spun at 12,000 g for 15 minutes. The top aqueous phase containing the RNA was mixed with pure isopropyl alcohol to precipitate RNA. RNA pellet was rinsed in 75% ethanol twice and then the pellet was dried and resuspended in nuclease-free water. The quantity and the quality of RNA was determined using ND1000 spectrophotometer (NanoDrop Technologies Wilmington, USA).

Equal amount of total RNA was converted to cDNA using iScript cDNA synthesis kit (Bio-Rad Laboratories, USA) using manufacturer’s guidelines. Quantitative real-time PCR was performed using a SYBR green based system obtained from Applied Biosystems, USA. PCR was carried out using the following primer sequences; VCAM-1 sense: 5’CAGGAGACATGGTGCTAAAG 3’, anti-sense: 5’CCAAGGAGGATGCAAAGTAG 3’, ICAM-1 sense: 5’ CTGGAGAGCACAAACAGCAGAG 3’and anti-sense: 5’ AAGGCCGCAGAGCAAAAGAAGC 3’. GAPDH (sense: 5’ ATGATTCTACCCACGGCAAG 3’ and anti-sense: 5’ CTGGAAGATGGTGATGGGTT 3’) with was used as the housekeeping gene. Quality check to ensure a single PCR product was carried out using the melting curve analysis. Double normalization was carried out with respect to total RNA and the housekeeping gene and the relative gene expression levels of the target genes were reported using the ΔΔCT method of analysis.

### 2.7 Immunostaining

Cells were fixed using 4% paraformaldehyde followed by 0.1% Triton permeabilization. Following this, the cell samples were blocked with 1% BSA solution for 1 hour and washed extensively. Cells were incubated in primary antibody (Santa Cruz, USA) overnight followed by 1 hour long incubation with secondary antibody tagged with TRITC and viewed under the fluorescent microscope.

### 2.8 Scanning Electron Microscopy Imaging

Fenestrations on the LSEC surface were assessed using a scanning electron microscope (SEM) (S-3000N, Hitachi, Japan). To prepare the samples for viewing, they were rinsed with PBS and fixed with 2.5% glutaraldehyde prepared in a solution of 0.1M sucrose in 0.1M sodium cacodylate buffer. After fixation, the samples were stored in cacodylate buffer and subjected to critical point drying using ethanol solutions of increasing strengths. At the final step of 100% ethanol treatment, the ethanol solution was removed using hexamethyl disilazane (HMDS) (Sigma Aldrich, USA) and the samples were allowed to air dry overnight. Following this the samples were sputter coated with gold-palladium and viewed using the SEM.

### 2.9 HA endocytosis assay

Cells were plated and allowed to recover for at least 2 hrs. Endocytosis media comprising of DMEM + 0.05% BSA supplemented with (a) 2 µg/ml ^125^I-HA (hot) and (b) 2 µg/ml ^125^I-HA + 100 µg/ml unlabeled HA (hot +cold) were freshly prepared. To prepare for the assay, LSEC media was aspirated and replaced with 300 µl of hot or hot +cold endocytosis media. Hot +cold media treatment served as the competitive internalization control for the assay to measure the specific uptake of iodinated HA. The cells were incubated at 37 ° C for 2 hours followed by gentle washing with Hank’s Buffered Salt Solution (HBSS) three times. The time point for the fresh LSECs was carried out after incubating the cells on “soft” or “stiff” substrate for 4 hours before adding endocytosis media. Similarly, for the 48 hours-readout, endocytosis media addition was carried out after cells were incubated on the PDMS substrates for 48 hours. The cells were lysed in 75 µl RIPA lysis buffer containing 0.5% NP-40 and the radioactivity was measured using a Gamma Scintillation Counter (Wallac 1470 (Perkin Elmer), Location). Scintillation count was considered directly proportional to the amount of ^125^I-HA uptake by the cells. HA uptake was normalized to total protein content of each sample, as determined by BCA colorimetric assay (Fisher Scientific, PA) and the results were expressed in terms of counts per minute/µg protein or CPM/µg protein.

### 2.10 Statistical Analysis

Data were expressed as the mean ± SD from three independent experiments. The difference between experimental groups was assessed using a one way ANOVA using the statistical feature available through Sigma Plot. Tukey test was applied to calculate the statistical significance and p < 0.05 was considered the threshold. Q tests were employed in order to identify the outliers among datasets.

## 3. Results

### PDMS substrates demonstrate uniform surface ligand density and LSEC attachment

We created *in vitro* 2D models representative of the mechanical microenvironment of the healthy and fibrotic liver using PDMS as the substrate. Tunability in the elastic modulus of the substrate was achieved by varying the weight based ratios of Sylgard 184 and Sylgard 527 (23). Precise weight ratios of Sylgard 184 to Sylgard 527 to obtain resultant substrates of physiologically relevant elastic moduli were empirically determined and the mathematical correlation between the varying amount of Sylgard 184 and Elastic Modulus of the substrate is demonstrated in **Fig 1**. The stiffness ratio of Sylgard 184:527 of 0:100 was used as the “soft” substrate with an elastic modulus of 2.3 ± 0.04 kPa, as determined through compression test. Similarly, Sylgard ratio of 15:85 was employed as the “stiff” substrate with a resultant elastic modulus of 55 ± 2.1 kPa **(Table.1).**

**Figure 1.**
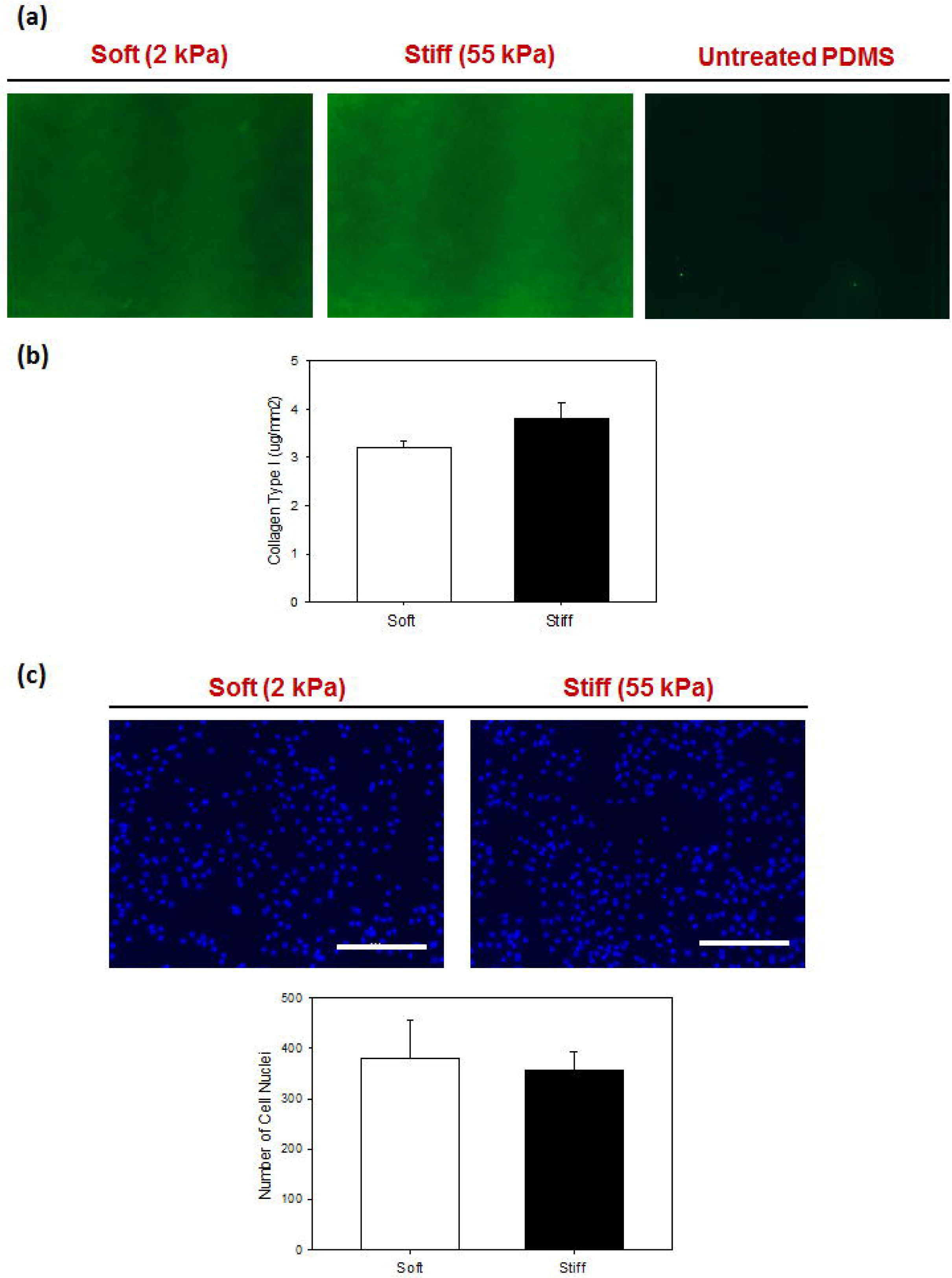
a) Collagen coating visualized of PDMS substrates using anti-collagen antibody mediated FITC staining (b) Quantification of collagen on the PDMS substrates using BCA assay and (c) Cellular attachment profile of LSECs on the soft and stiff substrates visualized using DAPI nuclear staining. Scale Bar = 50 microns

We subjected “soft” and “stiff” PDMS substrates to plasma treatment in order to functionalize the surface to allow for collagen type-1 binding. We used fluorescence microscopy in order to qualitatively test the efficacy of plasma treatment for functionalizing the surface and supporting uniform collagen coating. Both PDMS substrates coated with collagen FITC demonstrated a homogeneous green fluorescent signal on the surface as seen in **Fig. 1(a)**. The absence of green fluorescence signal in the untreated panel of **Fig. 1(a)** validates the necessity of oxygen plasma treatment in functionalizing the substrate surfaces. For a quantitative approach, we used BCA colorimetric protein assay to determine the amount of collagen deposited per unit surface area. As seen in **Fig. 1(b)**, soft and stiff substrates supported deposition of 3.2 ± 0.14 µg/mm^2^ and 3.7 ± 0.31 µg/mm^2^ collagen and as determined statistically, the difference in these values was found to be insignificant. (P > 0.05).

The ultimate goal of uniform collagen coating is to allow for uniform adhesion of primary LSECs. To assess uniform cellular adhesion, primary LSECs were seeded onto “soft” and “stiff” substrates and after allowing six hours of adhesion/recovery time, the cells were fixed with paraformaldehyde and nuclei stained with DAPI. Assuming a direct correlation between the number of nuclei and the cell number for LSECs (mononuclear cell type), we observed that the cells demonstrated a uniform monolayer-like cell distribution, as seen in **Fig. 1 (c)** and quantitative analysis of nuclei per unit area also corroborated the similarity in cell number between the two substrates.

### Substrate stiffness mediates change in LSEC morphology and cytoskeletal organization

After confirming the occurrence of similar ligand density and cell adhesion profile, we analyzed the role of mechanical stiffness in the regulation of cellular phenotype considering cell morphology as a preliminary marker. We found that after 48 hours, LSECs on “soft” substrate had the typical “fried egg” morphology that is reported in literature when LSECs are cultured on 2D surfaces, as opposed to LSECs cultured on “stiff” substrate, which appeared to have developed elongated “fibroblast-like” morphology **(Fig. 2(a))**. Additionally, LSECs on “stiff” substrate demonstrated heterogeneity regarding cellular morphology and cellular area when compared with LSECs on “soft” substrate.

**Figure 2.**
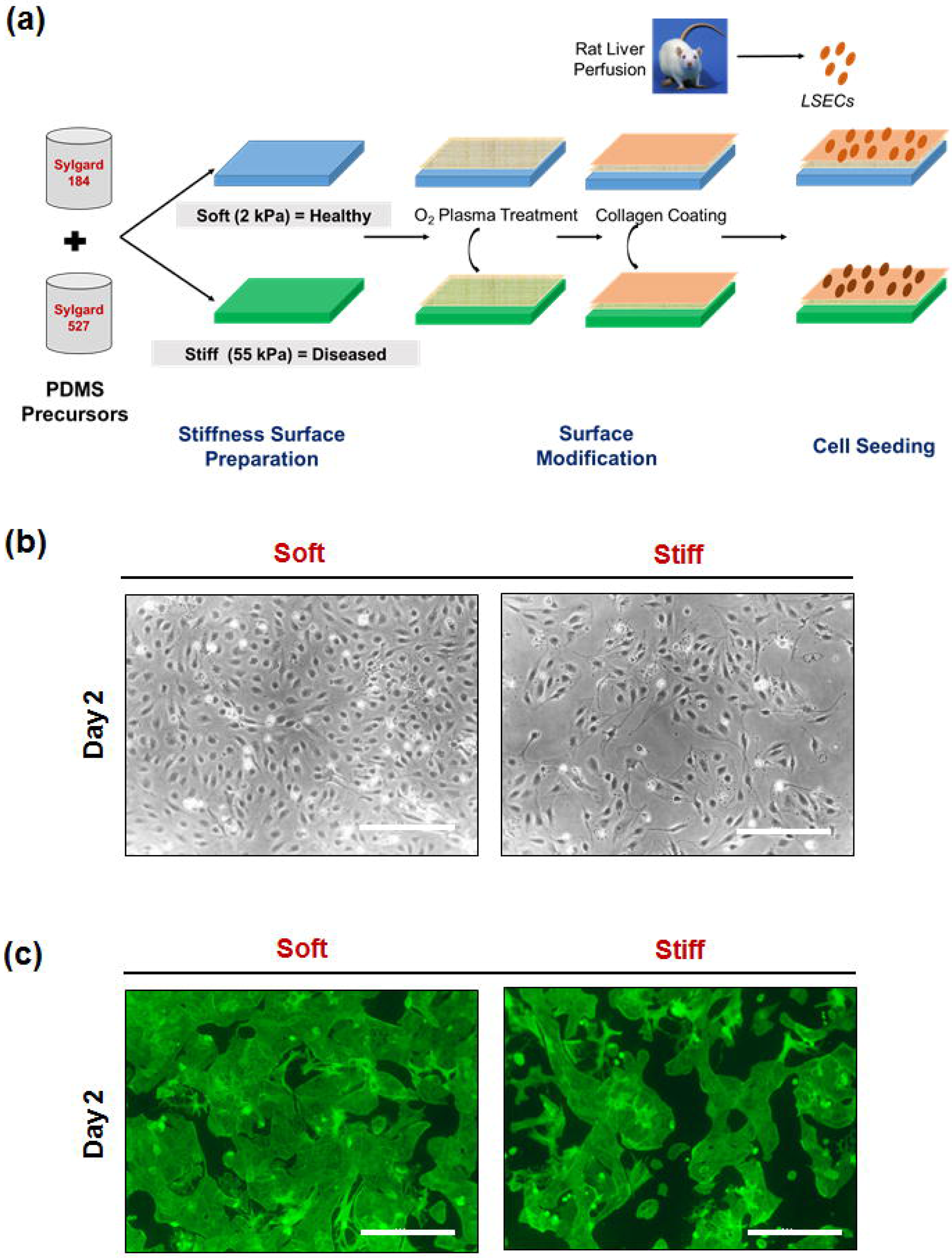
(a) Schematic for the design of *in vitro* models that represent healthy and diseased liver microenvironment (b) phase contrast imaging of LSECs morphology and (c) F-actin-phalloidin-Alexa 488 staining of LSECs to visualize the cell morphology and stress fibers. Scale bar = 50 microns

Since altered mechanical stimulus is synonymous with cytoskeletal rearrangement, we stained the cells with Phalloidin-FITC to visualize the F-actin filaments. LSECs cultured on “soft” substrate demonstrated an overall a lower expression of F-actin fibers which also appeared dispersed throughout the cell body, as opposed to cells on “stiff” substrate, where the appearance of actin stress fibers was prominent. Additionally, LSECs on “stiff” substrate also demonstrated the tendency to form clusters **(Fig. 2(c))**. Collectively, the morphological observations on LSECs suggest that “soft” substrate maintained the cells in a differentiation phenotype, as opposed to the “stiff’ substrate.

### Fibrotic substrate stiffness results in defenestration of LSECs

LSEC fenestrations are among the most extensively characterized feature of these cells and, loss in fenestrations is a gold standard marker for capillarization/dedifferentiation. We employed an SEM based investigation to study the mechanical stiffness based alteration in LSEC fenestration size distribution and frequency in the healthy and the fibrotic liver. As seen in **Fig. 3**, LSECs cultured on “soft” substrate for 24 hours demonstrated the characteristic presence of fenestrations that were clustered into the typical sieve-plate like structures. There were some isolated/ungrouped fenestrations which could potentially be representative of fragments of broken sieve plates. LSECs cultured on “soft” substrate for a longer duration of 48 hours still displayed the intact maintenance of fenestrations that were grouped in sieve plates. On the contrary, LSECs on “stiff” substrate demonstrated complete defenestration as early as 24 hours and demonstrated the same phenomenon when visualized at 48 hours. These observations strongly suggest the role of mechanical stiffness in triggering a loss in fenestration and capillarization in LSECs. The frequency and size distribution of fenestrae are documented in **Table. 2**. LSECs cultured on “soft” substrate had comparable fenestration sizes and density, at 24 hours and 48 hours in culture. We observed that fenestrations were 119 ± 45 nm at the density of 2-4 per µm^2^.on Day 1 and 128 ± 52 nm in size at the density of 2-5 per µm^2^.on Day 2.

**Figure 3.**
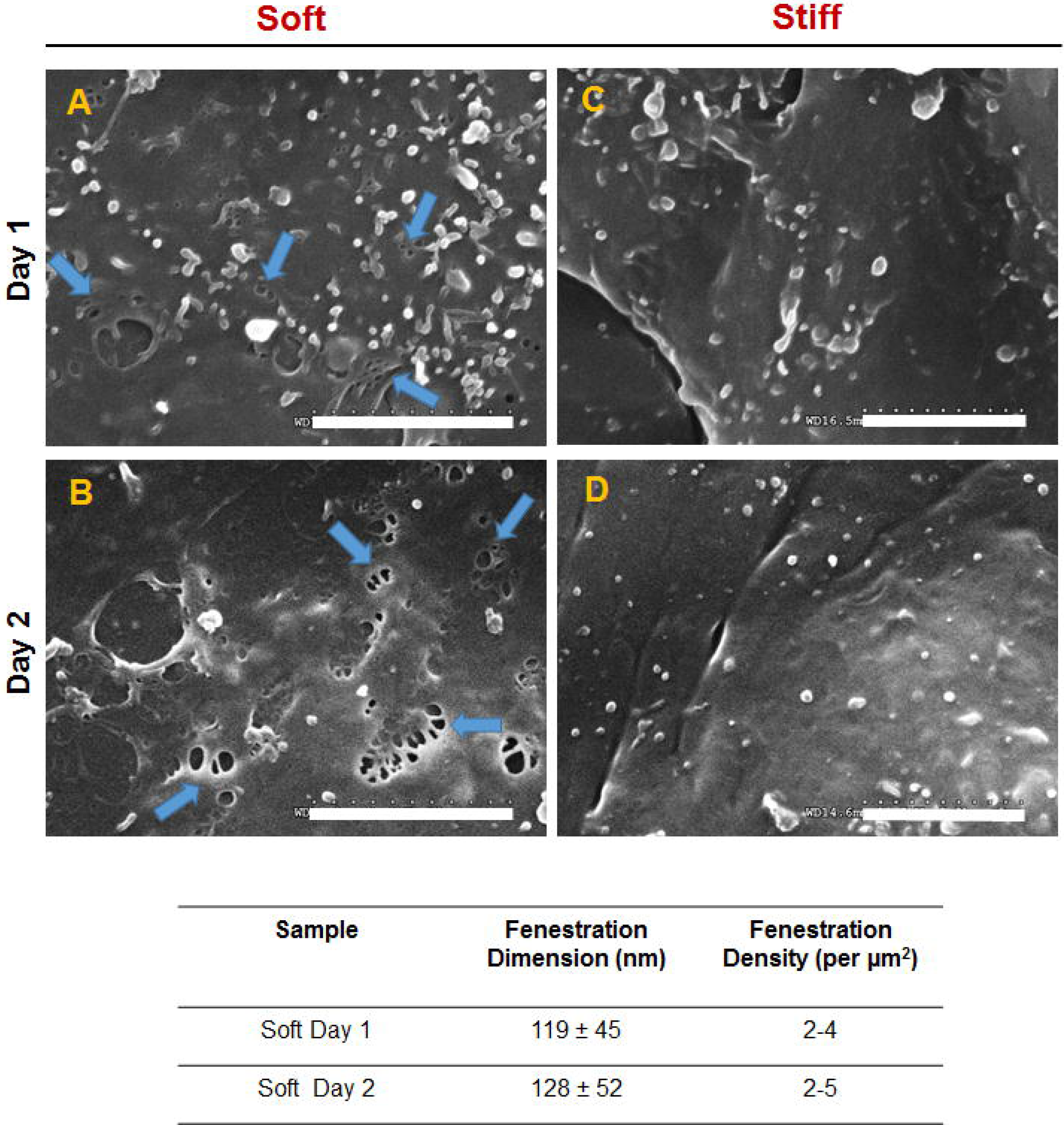
(a) SEM images of the fenestrations on LSECs after 24 hours in culture. A-B represent LSECs cultured on the soft substrate and C-D represent LSECs on the stiff substrate. Blue arrows point to the fenestrations. Scale bar = 5 microns

**Table 2.** Fenestration dimension and occurrence per unit area of LSECs cultured on the “soft” substrate for 24 and 48 hours.

### Fibrotic substrate stiffness results in higher expression of cell adhesion molecules VCAM-1 and ICAM-1 in LSEC

Capillarized LSECs experience a phenotypic drift towards general vascular endothelial cell type in the liver during liver disease and express altered levels of cell adhesion molecules that alters the inflammatory response of the endothelium. We probed for the expression levels of vascular cell adhesion molecule (VCAM-1), and intercellular cell adhesion molecule (ICAM-1) in LSECs on “soft” and “stiff “substrates. As seen in **Fig. 4 (a)**, the gene expression analysis shows that when cultured on ‘stiff” substrate for 48 hours, the expression level of VCAM-1 and ICAM-1 in LSECs increases significantly. When normalized with respect to LSECs cultured on “soft” substrate for analysis purpose, the expression of VCAM-1 is 1.53 ± 0.533 folds higher, and ICAM-1 is 2.1 ± 0.7 folds higher than the “soft” substrate. We observed a similar pattern in the protein expression analysis of LSECs as well. (Fig. 4 (b)). Western blotting followed by densitometric analysis for VCAM-1 and ICAM-1 protein levels demonstrated that the LSECs cultured on “stiff” substrate had a higher expression level, as compared to “soft” substrate. These results are in agreement with the defenestration data and collectively suggest that LSECs cultured on the fibrotic liver mimic demonstrate a capillarized phenotype.

**Figure 4.**
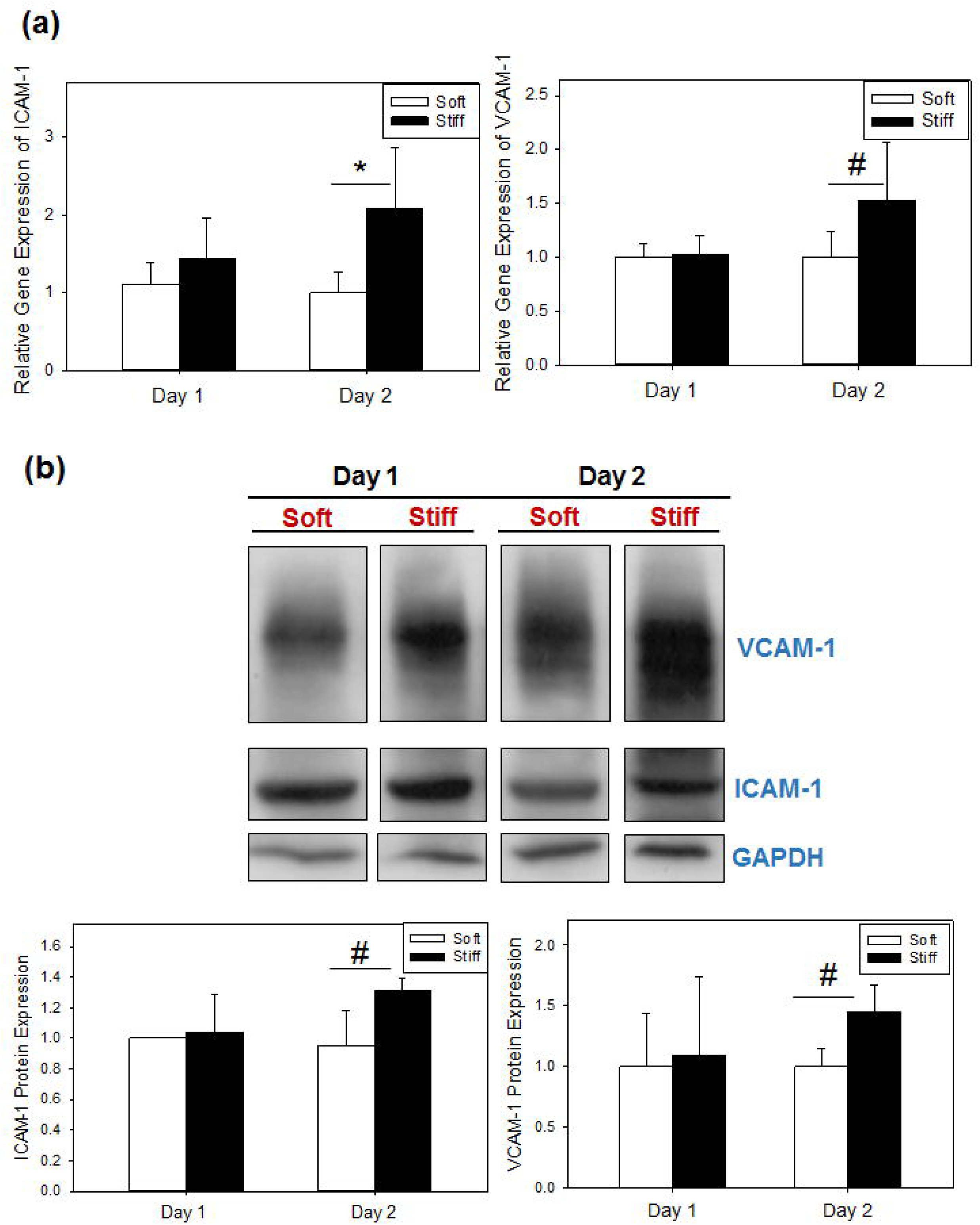
(a) Gene expression analysis of VCAM-1 and ICAM-1 using qPCR (b) Western blotting analysis of the expression of ICAM-1 and VCAM-1 in LSECs cultured on soft and stiff substrates and densitometric plot for western blot (to the total protein content # : p value < 0.05 and * p value < 0.005

### Fibrotic substrate stiffness mediates loss in HA endocytosis function in LSECs

LSECs constitute an active scavenging system that regulates the availability of different ligand types in the lumen of the liver. Several different scavenger receptors are involved in this scavenging process, including hyaluronan receptors. We evaluated the specific clearance of hyaluronic acid (HA) of LSECs on “soft” and “stiff” substrates. As seen in **Fig. 5(a)**, fresh LSECs that were seeded onto “soft” and ‘stiff” substrate, followed by 4 hours of recovery time before evaluating their HA endocytosis activity, demonstrated a similar functional maintenance with LSECs on “soft” at 49.2 ± 4.2 cpm/ µg total protein and ‘stiff” at 41.3 ± 8.6 cpm/ µg total protein. On the contrary, when the cells were subjected to different mechanical microenvironment for duration of 48 hours, the endocytosis functional maintenance appeared substantially different. LSECs on “soft” substrate endocytosis was around 26.3 ± 7.6 cpm/ µg total protein, whereas, LSECs on “stiff” substrate demonstrated less than 50% of the “soft” endocytosis capacity at 12 ± 0.6 cpm/ µg total protein.

**Figure 5.**
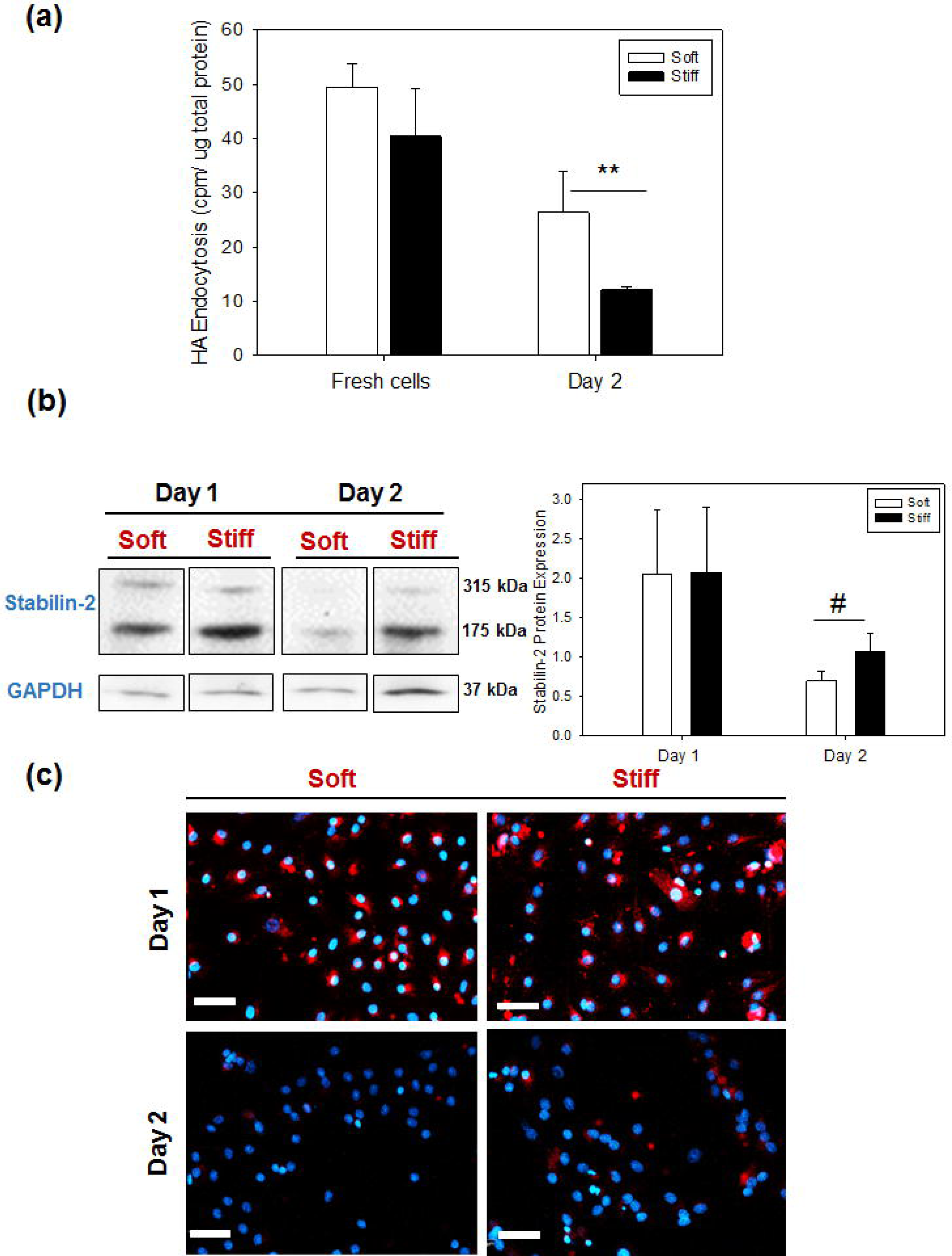
(a) Quantification of HA endocytosis function normalized to the total protein content, # : p value < 0.05 (b) Western blotting analysis of the expression of Stabilin-2 in LSECs cultured on soft and stiff substrates and densitometric plot for stabilin-2 western blot # : p value < 0.05 and (c) Immunostaining = blue represents DAPI stained nuclei and Red represents stabilin-2, scale = 40 microns

The uptake of HA in LSECs is specifically carried out by hyaluronan receptor stabilin-2 (HARE protein). We evaluated the protein expression levels of stabilin-2 to establish a correlation with the observed endocytosis activity. As seen in **Fig. 5(b)**, at 48 hours in culture, LSECs on “stiff” substrate demonstrated a higher expression of Stabilin-2, as compared with LSECs on “soft” substrate. These results were also corroborated in the immunostaining of LSECs for stabilin-2, as seen in **Fig. 5(c)**. Higher expression of stabilin-2 in “stiff” substrate accompanied by a lower HA endocytosis function might suggest that stabilin-2, even though is abundant in the capillarized LSECs, is not functional in nature. Detailed molecular investigations pertaining to protein structure and function will be required to establish the mechanism contributing to the loss in stabilin-2 functionality.

## 4. Discussion

LSECs are the second most abundant hepatic cell type, and their unique functional properties are vital for the maintenance of homeostasis (1). These cells act as molecular sieves owing to their fenestrated surface, play an active role in scavenging of molecules through endocytosis, and, regulate the phenotypic stability of other parenchymal and non-parenchymal cells through paracrine signaling (3). Conversely, endothelial dysfunction results in several hepatic complications, including liver fibrosis. DeLeve and coworkers demonstrated that capillarized LSECs trigger activation of hepatic stellate cells resulting in a cascade of pro-fibrotic events in the liver and this phenomenon was found to be reversible if LSECs phenotype was restored (25). Capillarization of LSECs also directly affects the communication between hepatocytes and LSECs, thereby mediating loss in hepatic functions. Realization of the pertinent role of LSECs in liver fibrosis progression has several clinical implications. Establishing the mechanistic aspects of LSECs phenotypic deterioration can result in the identification of novel biomarkers that could support early detection of liver fibrosis and thereby increase the treatment efficacy. Additionally, this research would also enable development of anti-fibrosis strategies that target restoration of LSECs capillarization.

LSECs were identified as a specialized hepatic cell-type, as opposed to being passive barrier cells, through the seminal work by Wisse and group involving extensive SEM analysis and since then, many important functional and structural aspects of these cells have been uncovered (26). Despite the progress, LSEC biology has been a challenging field of research since the standard vascular endothelial cell culture techniques fail to support LSECs *in vitro* and result in cells undergoing spontaneous dedifferentiation (27) (28). Most *in vitro* models rely on the culture of LSECs on protein coated polystyrene dishes, which result in these cells undergoing a rapid loss in cellular fenestrations in less than 48 hours. Several microenvironmental elements contribute to the functional maintenance of LSECs. To our best knowledge, only two studies have analyzed the role of mechanical microenvironment on LSEC phenotypic characterization. Juin and coworkers utilized polyacrylamide gel-based platform to show that LSECs suffered loss in podosome formation when subjected to increasing substrate rigidity (22). Ford and coworkers utilized collagen gel substrates to demonstrate the loss of fenestrae in LSECs as a result of increased elastic modulus (21). Our study is a novel attempt at understanding the nature of mechanosensitivity in LSECs as our model is the first to recreate the 2D mechanical microenvironment of the healthy and fibrotic liver without altering the surface ligand density. Additionally, we have also reported the loss in the functional aspects of LSECs mediated by changing mechanical microenvironment.

Sylgard 184 has been extensively employed in biomedical applications as a result of its superior biocompatibility, optical transparency, and cost-effectiveness (29). On the contrary, Sylgard 527 is a relatively less popular substrate in terms of cell-related applications. Our group has previously succeeded in engineering *in vitro* mechanical model to study the phenotypic changes in hepatocytes as observed in healthy and fibrotic livers (19), and this study has established the efficacy of utilizing the blended mixture of Sylgard 184 and 527 without resulting in undesired cellular effects. The blending of Sylgard 184 and 527 to create tunable substrate stiffness is a highly versatile and reproducible technique that yields substrates of physiologically relevant mechanical stiffness range corresponding to a healthy liver tissue and advanced stage liver fibrosis. Another salient feature of the PDMS-based model is the ability to uncouple the varying stiffness of the substrate from ligand density changes. The primary disadvantage of employing bio-responsive substrates such as collagen gel, heparin or fibrin gel to create mechanical models of tissues is that to increase the elastic modulus of the substrate, the inherent ligand composition has to be increased, which results in a dissimilar amount of bio-responsive ligand availability for cells on the surface. This phenomenon could potentially introduce bias in the interpretation of the mechanosensitive behavior of the cells in question.

Our model is efficient in supporting ideal cellular adhesion and maintenance, which is a crucial factor in the study of LSECs, as they are primary cells and can be difficult to maintain *in vitro*. LSECs cultured on “soft” substrate depicted the typical fried egg morphology for up to 48 hours which was lost in LSECs on “stiff” substrate. Similarly, LSECs cultured on “stiff” substrate developed substantially higher actin stress fibers **(Fig. 2(c))**. These morphological changes establish the mechanosensitivity of LSECs as changes in cytoskeletal framework and increase in actin stress fibers are observations commonly in agreement with other adherent cell-types subjected to mechanical stress (30). In the context of LSECs, cytoskeletal reorganization has special repercussions on the maintenance of fenestrations.

Due to their lack in basement membrane, presence of fenestration in LSECs is a critical morphological feature to regulate size based transport of entities to the Space of Disse thereby gaining access to hepatic parenchyma (31). Changes in fenestration frequency, size and spatial arrangement are correlated with the progression of several liver pathophysiological conditions and disappearance of fenestrations is considered analogous to capillarization (1). Our study suggests that in the event of liver fibrosis, the altered mechanical environment is a driving force in the loss of fenestration in LSECs.

Cell-matrix interactions in all endothelial cells are governed by adhesion proteins such as the vascular cell adhesion molecule (VCAM) and the intercellular cell adhesion molecule (ICAM) (32). In a healthy liver, LSECs express a basal level of ICAM-1, but in case of liver injury due to viral infections or chronic inflammatory conditions, the expression level of ICAM-1 goes up, indicating the phenotypic adaptation that LSECs undergo, as a response to changing microenvironment (33). Similarly, a study using rodent model for liver cirrhosis demonstrated that as compared to the healthy sinusoid, cirrhotic sinusoid had a substantially higher expression of VCAM-1 (34). Physiologically, higher expression of adhesion molecule represents elevated inflammatory response, mediated by adhesion and transmigration of lymphocytes and macrophages. Our study indicates that LSECs cultured on “stiff” substrate and consequentially exposed to fibrosis-like microenvironment, express a higher level of VCAM-1 and ICAM-1, depicted at both mRNA and protein level **(Fig. 4)**. This result suggest that the mechanical stress causes a pro-fibrotic phenotypic change in LSECs. Collectively, loss in fenestration and higher expression of VCAM-1 and ICAM-1 in LSECs cultured on the in vitro model of fibrosis suggest the substantial role of mechanical stiffness in exacerbating capillarization of these cells. Our model could be a valuable tool towards the short term study of capillarized LSECs in order to investigate the mechanistic aspects associated with capillarization phenomenon and their implications on LSEC functions.

LSECs function as a scavenging system allows for recycling of macromolecular waste from the lumen such as ECM remnants, metabolic end products, lipoproteins and viral particles (35) (36) (37) (38). Hyaluronic acid (HA) is a polysaccharide that forms a major constituent of the ECM and removal of excess or damaged HA is vital for the maintenance of blood serum viscosity (39). In the liver, HA clearance is carried out exclusively by stabilin-2, a large type 1 membrane protein containing of isoforms of 175kDa and 315kDa (40). Serum HA levels have been used as an indicator for several liver metabolic diseases (8). Our *in vitro* fibrosis mimic (LSECs on the stiff substrate) reveals that LSECs undergo a loss in HA endocytosis function which suggests that along with the increase in HA production in a fibrotic liver, the increase in serum HA level could be due to the loss of endocytosis function in LSECs. The study reported here is the first to show a direct correlation between the functional aspects of LSECs and the mechanical microenvironment of the liver. The loss in HA endocytosis ability is observed despite there being an upregulation in stabilin-2 expression. This observation could potentially mean that even though the stabilin-2 expression is not downregulated, the protein exists in a non-functional form.

Overall, previous research establishes that the appropriate microenvironment mimicking can prolong the survival of LSECs *in vitro* (41)(27, 42). From the perspective of liver tissue engineering, our study illustrates that along with the chemical/ligand composition of the culture microenvironment, the mechanical aspects of the culture substrate are of prime importance while developing *in vitro* models that can sustain the differentiated phenotype of LSECs for a longer duration. Our healthy liver mimic (soft substrate) demonstrates a successful maintenance of LSEC in their differentiated phenotype and conversely, the fibrosis mimic (stiff substrate) is a snapshot of the capillarized LSEC phenotype. Understanding the role of the various LME elements in regulating the function of LSECs is vital towards both functional hepatic tissue engineering and establishing *in vitro* models to study the various pathophysiological conditions. The models generated can be applied towards high throughput identification of LSECs biomarkers for liver fibrosis and also as an efficient preclinical drug screening platform to investigate anti-fibrosis therapies.

